# Mechanistic insights into the activation of lecithin-cholesterol acyltransferase in therapeutic nanodiscs composed of apolipoprotein A-I mimetic peptides and phospholipids

**DOI:** 10.1101/2022.06.09.495129

**Authors:** Laura Giorgi, Akseli Niemelä, Esa-Pekka Kumpula, Ossi Natri, Petteri Parkkila, Juha T. Huiskonen, Artturi Koivuniemi

## Abstract

The mechanistic details behind the activation of lecithin-cholesterol acyltransferase (LCAT) by apolipoprotein A-I (apoA-I) and its mimetic peptides are still enigmatic. Resolving the fundamental principles behind the LCAT activation will facilitate the design of advanced HDL-mimetic therapeutic nanodiscs for LCAT deficiencies and coronary heart disease, and for several targeted drug delivery applications. Here, we have combined coarse-grained molecular dynamics simulations with complementary experiments to gain mechanistic insight into how apoA-I mimetic peptide 22A and its variants attune LCAT activity in peptide-lipid nanodiscs. Results highlight that peptide 22A forms transient antiparallel dimers in the rim of nanodiscs. The dimerization tendency considerably decreases with the removal of C-terminal lysine K22, which has also been shown to reduce the cholesterol esterification activity of LCAT. In addition, our simulations revealed that LCAT prefers to localize to the rim of nanodiscs in a manner that shields the membrane-binding domain (MBD), αA-αA’, and the lid amino acids from the water phase, following the previous experimental evidence. Meanwhile, the location and conformation of LCAT in the rim of nanodisc are spatially more restricted when the active site covering lid of LCAT is in the open form. The average location and spatial dimensions of LCAT in its open form were highly compatible with the electron microscopy images. All peptide 22A variants studied here had a specific interaction site in the open LCAT structure flanked by the lid and MBD domain. The bound peptides showed different tendencies to form antiparallel dimers and, interestingly, the temporal binding site occupancies of the peptide variants affected their *in vitro* ability to promote LCAT-mediated cholesterol esterification.

## Introduction

High-density lipoprotein (HDL)-mimetic nanodiscs (or synthetic HDL, sHDL) are promising therapeutic agents in treating lecithin-cholesterol acyltransferase (LCAT) deficiencies and coronary heart disease (CHD) (1,2). Moreover, because of their unique intrinsic targeting capabilities, sHDLs have been demonstrated to serve as versatile molecular platforms in several drug delivery applications, including the presentation of cancer neoantigens to the immune system, targeted delivery of liver X receptor agonists to arterial macrophages, and the delivery of cytotoxic cancer therapeutics through overexpressed SR-BI receptors (3–5). Therefore, many research studies have been recently carried out to understand their genesis, structural, mechanistic, and pharmacokinetic properties to tailor advanced medicinal nanodiscs for different therapeutic areas (6–15).

HDL-mimetic nanodiscs are comprised of different lipids and apolipoprotein A-I (apoA-I) proteins, or peptides mimicking the lipid-binding characteristics of apoA-I (∼20 aa in length). The production of nanodiscs composed of apoA-I mimetic peptides and phospholipids (PLs) is relatively straightforward *in vitro* exploiting, for example, different sonication and freeze-thaw techniques (16). Molecular dynamics (MD) simulations and experimental evidence have shown that in the nanodisc the lipids form a bilayer/bicelle-like structure where the amphiphilic α-helical peptides reside in the rim of the disc, shielding the hydrophobic acyl chains of PLs from aqueous surroundings and stabilizing the nanoparticles (11,17–19). Depending on the type of apoA-I mimetic peptides and PLs, and their ratio, the overall size of nanodiscs ranges from a few to tens of nanometers based on dynamic light scattering (DLS) and electron microscopy (EM) analyses (10,20).

The chief idea and hypothesis behind the use of apoA-I mimetic peptides and their lipid complexes in treating CHD in humans are backed up by numerous animal studies showing that they possess various atheroprotective effects (1). Clinical studies, however, suggest that the injection of apoA-I-based HDL-mimetics to patients with a recent history of acute coronary complications may be an unviable strategy, although the largest of the trials, AEGIS-II, is still proceeding (21). Therefore, in the future, longer interventions starting already at the early indications of CHD-relevant metabolic disorders might be more indicative of the beneficence of sHDL nanoparticles in alleviating the complications of atherosclerosis. In this context, peptide-based synthetic HDL-mimetic nanodiscs have been designed and investigated as they are less costly and less laborious to manufacture on a large scale compared to the complex extraction, purification, and homogenization process of full-length apoA-I or bloodborne native HDL particles.

As in the case of native HDL particles, research suggests that efficient cholesterol (CHOL) efflux from cells, and removal of surplus CHOL, is an essential therapeutic feature of synthethic HDL-mimetic particles (22,23). Therefore, this has long been a sought-after property of synthetic HDL-mimetic nanodiscs and apoA-I mimetic peptides. The native-like maturation of the nanodiscs by lecithin-cholesterol acyltransferase (LCAT) has been regarded as a desired feature as it would lead to the esterification of free CHOL and the formation of hydrophobic cholesteryl ester(CE)-rich core. The CHOL esterification mediated by LCAT is believed to augment the CHOL loading capacity of HDL particles and synthetic nanodiscs, increasing the free CHOL gradient between HDL-mimetics and arterial macrophages. This, in turn, increases the efficiency of reverse cholesterol transport (RCT) (24). However, it remains unclear if LCAT activity can be targeted in a manner that is beneficial in the treatment of CHD through RCT. This is partly because of the much faster turnover of free CHOL in HDL when compared to LCAT-mediated esterification rate and the capacity of LCAT to esterify CHOL in very low density and low-density lipoproteins (VLDL and LDL), so-called β-LCAT activity (25–28). Nevertheless, various LCAT-targeted therapeutics are in development against LCAT deficiencies and cardiovascular diseases (29,30). To this end, several apoA-I mimetic peptides have been designed, and, interestingly, some of them, such as 22A (Esperion Therapeutics, ESP24218), activate LCAT nearly as well as native apoA-I (31). Yet, the mechanistic principles of how apoA-I mimetic peptides can activate LCAT are poorly understood. Revealing the cofactor mechanism of apoA-I mimetics and features affecting it will open new avenues, for example to improve the targeting, pharmacokinetic, and –dynamic properties of apoA-I mimetic peptides in a manner that makes them more fitting for long-term medication and clinical trials.

Previous functional studies carried out with peptide 22A nanodiscs have indicated that positively charged C-terminal lysine K22 is essential in the CHOL esterification since it has been found that the removal of K22 decreases the CHOL esterification down by ∼60 percent (6). Further, it has been shown that 22A undergoes rapid hydrolysis in plasma to form 21A peptides without the C-terminal K22 potentially hindering the use of 22A peptides, for example to treat LCAT deficiencies (6). However, the removal of C-terminal K22 has shown earlier no effect on phospholipolytic action mediated by LCAT. (6) The latter finding agrees with the research conducted by Gorshkova et al. showing that the mutation of the positively charged amino acid R123 in native apoA-I structure with a 5/5 registry affects only CHOL esterification but not the hydrolysis of phospholipids (32). In addition, Cooke et al. have pointed out the importance of apoA-I registries in native HDL particles by demonstrating that the activity of LCAT is regulated by a thumbwheel mechanism in which the correct positioning of amphiphilic apoA-I helixes is crucial for the full LCAT activity (33). Very recently, the location of LCAT in discoidal HDL particles has been resolved with negative stain EM. This has revealed that LCAT prefers to bind to the edge of discoidal HDL particles and next to the helixes 5 and 6 of apoA-I (34). These data also suggest that helixes 5 and 6 create a path from the lipid bilayer to the active site of LCAT. Based on these findings alone, it is intriguing that such small peptides can activate LCAT with the same potency as apoA-I.

Mechanistic level understanding is required not only to tune the targeting properties of therapeutic nanodiscs but it would also provide insights into how apoA-I activates LCAT. However, the structural organization of 22A peptides and how they interact with LCAT in the peptide-lipid nanodiscs remain poorly understood. In particular, it is unclear how different apoA-I mimetic peptides are structurally organized in nanodiscs and if this plays a significant role in the activation of LCAT? Further, the knowledge regarding the principal binding site and interaction mode of apoA-I mimetic peptides in the structure of LCAT is missing. Most importantly, it is elusive if a similar kind of dimer arrangement of apoA-I mimetic peptides is required for LCAT activation as in the case of apoA-I.

In this study, we carried out coarse-grained MD simulations and complementary experiments aiming to characterize the structural and dynamical properties of apoA-I mimetic peptides in therapeutic nanodiscs with and without LCAT. Since the relative helical position of two apoA-I monomers in HDL affects LCAT activation potency, we hypothesized that the dimerization of 22A peptides and their relative orientation to each other would play a significant role in LCAT activation. Further, we investigated the location, orientation, and positioning of LCAT with respect to the nanodisc surface as it alone might regulate the access of PLs and CHOL into the active site of LCAT. Moreover, although LCAT prefers to interact with apoA-I in the rim of HDL particles, we wanted to examine if this also holds in nanodiscs composed of apoA-I mimetic peptides and PLs. We also sought to find specific interaction sites for apoA-I mimetic peptides in the lipid-LCAT interface and the effect of 22A variants on all the above features.

Our results highlight that peptide 22A forms antiparallel dimers in the rim of nanodiscs and that the removal of C-terminal lysine K22, shown to reduce the CHOL esterification activity of LCAT, abolishes this dimerization. Molecular dynamics simulations with LCAT revealed that the enzyme prefers to localize to the rim of nanodiscs and, although the conformation of LCAT is dynamic, it adopts a spatial configuration that was highly compatible with the negative staining EM images. Further, all peptide 22A variants studied here had a specific interaction site in the open LCAT structure flanked by the lid and membrane-binding domain (MBD) which was coined as site A. While residing in site A, the peptides showed different tendencies to form antiparallel dimers and the temporal occupancies of the peptides in the site strongly followed their potencies to activate LCAT *in vitro*. Mechanistic insights provided here open up avenues to design pharmacologically more potent apoA-I mimetics and therapeutic nanodiscs for different medicinal applications.

## Methods

### Molecular dynamics simulations

To examine how the apoA-I mimetic peptides behave on sHDL nanodiscs, coarse-grained molecular dynamics simulations were conducted with the GROMACS simulation package (35) using the Martini 3.0 force field (36). All programs preceded by gmx described later are a part of this package. The parameters for 1,2-dimyristoyl-sn-glycerol-3-phosphocholine (DMPC) were obtained from the Martini website (cgmartini.nl), and the parameters for the peptides and LCAT were generated via the Martinize2 program. MD simulations were run with a 20 fs timestep. Lennard-Jones interactions were handled with 1.1 nm cut-off with potential-shift-verlet vdw-modifier. Electrostatic interactions were handled using a reaction-field scheme with 1.1 nm cut-off. The dielectric constant was set to of 15. The systems were coupled to a velocity rescaling thermostat set to 320 K with a 1 ps coupling constant (37). The pressure was handled by an isotropic Parrinello-Rahman barostat (38) with pressure set to 1 bar with a 12 ps coupling constant and a 3×10^−4^ bar^-1^ compressibility.

Each simulated nanodisc was comprised of 200 DMPC molecules and 28 peptides. Simulated peptides and their sequences were: 22A (PVLDLFRELLNELLEALKQKLK), 22A-K ((PVLDLFRELLNELLEALKQKL), 22A-K22Q (PVLDLFRELLNELLEALKQKLQ) and 22A-R7Q (PVLDLFQELLNELLEALKQKLK). The secondary structures of peptides were set to α-helical. They were prepared by first placing the lipids and peptides randomly into a 10 × 10 × 10 nm empty box. Then they were minimized and ran for a few nanoseconds with the barostat turned on to compress the molecules together. Once compressed, the molecules were placed in a 20 × 20 × 20 nm box and solvated with 26000 Martini water model beads. After minimization, these systems were run up to 20 µs. As the discs formed rapidly and the number of contacts between lipids and peptides equilibrated in ∼100 ns, the whole trajectories were used in the analysis. This afforded four systems: 22A, 22A-K, 22A-R7Q, and 22A-K22Q. Once the nanodisc had formed, CG LCAT in an open or closed state via its membrane-binding domain was placed into the middle of the nanodiscs surface to the lipid-water interface. The coordinates for open LCAT were obtained from PDB code 6MVD and for closed LCAT from 4XWG. Before coarse-graining, 4XWQ structure residue Y31 was back mutated to C31. For all peptides, triplicate simulations of 10 µs (40 µs in effective Martini time) were run with both LCAT types. Before coarse-graining, 4XWQ structure residue Y31 was back mutated to C31.

First, simulation systems without LCAT were scrutinized. The nanodiscs were oriented so that the disc normal was parallel to the z-axis with gmx editconf. The radial distribution functions, peptide angle distributions relative to the z-axis, and the rotational autocorrelation functions of the peptides were calculated with gmx rdf, gmx gangle, and gmx rotacf, respectively. For analysing the orientations of the peptides, residue 1 and the last residue described each peptide. The dimerization tendency of the peptides was determined with gmx cluster and was investigated more carefully by plotting the angle with gmx gangle as a function of the distance between residue 13 backbone beads with gmx mindist for all peptide pairs. The plots were generated with the Matplotlib library (39) and the snapshots of the simulations were rendered with the VMD software (40).

Next, the generic behavior of LCAT was characterized. The number of contacts between LCAT backbone beads and peptide backbone beads as a function of time was calculated with gmx mindist using a distance cutoff of 3 nm. The LCAT beads were grouped, so the number of contacts is the number of peptide beads in the vicinity of the whole LCAT structure. These graphs were used to determine when the systems were in equilibrium (i.e. when LCAT assumed a stable pose on the perimeter of the discs). The pose of LCAT relative to that of the nanodisc was determined by a custom script. The systems were oriented as previously described, with the disc normal placed parallel to the Z-axis. The script determines the center of mass of the nanodisc (DMPC-COM) and calculates the distance of a backbone bead of a central residue of LCAT (CYS31) along the XY-plane and Z-axis separately to it. Then, a plane is defined by DMPC-COM, CYS31, and the z-axis. By determining the polar coordinates of the backbone bead of ILE326 relative to the plane and CYS31, the script attains the pitch and yaw angles of LCAT. If the CYS31 to ILE326 vector projected on the plane points orthogonally towards a z-axis line that passes DMPC-COM, the pitch angle is set to 0 degrees, and the angle increases up to 360 degrees as the vector rotates: the angle is 90 degrees when it points towards positive z, 180 degrees when it points orthogonally away from the line and 270 degrees when pointing towards negative z. Finally, the vector from CYS31 to the backbone bead of GLY308 is used to determine the roll of LCAT along CYS31 to ILE326. As orienting the nanodisc in each frame caused LCAT to be randomly pointed upside down, this angle (roll) was used to correct the orientation of LCAT relative to the nanodisc. After plotting the CYS31 XY-plane distance from DMPC-COM as a function of the z-axis position of each frame and coloring the points based on the pitch angle (denoted by θ), a clear correlation was noticed between the latter two parameters. The plots were created with Matplotlib (39) and the R^2^ value of the linear correlation between the z-axis position and the pitch angle was calculated with the SciPy library (41).

To see which LCAT residues were primarily in contact with the 22A based nanodisc, average solvent-accessible surface areas (SASAs) per residue were calculated with gmx sasa. The average results of the triplicate simulations are reported. The van der Waals radii were set to match Martini beads (regular = 0.264 nm, small = 0.225 nm, and tiny = 0.185 nm). The probe radius was set to 0.264 nm (regular bead). Supplementary simulations of CG LCAT in an open or closed state in a pure water system were run for 200 ns for reference. A comparison between the SASAs of LCAT open on a disc and the SASAs of LCAT closed in water was also performed.

Finally, peptide behavior relative to LCAT was inspected. The trajectories were rotationally and translationally fitted to the backbone beads of LCAT, and spatial density maps for the peptide beads were formed with gmx spatial using a bin width of 0.2 nm. Upon visualization we noted that in open LCAT simulations peptides had a tendency to stay in a groove defined by the lid and the MBD.

Another custom script was written to characterize these peptides. First, the script finds the peptide with the minimum combined distance between peptide-PRO1-LCAT-PRO232 and peptide-LEU21-LCAT-TRP48. If the PRO1-PRO232 distance is less than 1 nm and LEU21-TRP48 less than 2 nm, the peptide was considered to be located in the binding site. The number of frames when a peptide was bound divided by the total number of frames gives the percentage of time a peptide occupied the binding site. As these results did not converge between triplicate simulations of 10 µs, the occupancy percentages were calculated again after running the open LCAT simulations until 20 µs (80 µs effective Martini time). Additional data was gathered on the behavior of these binding events (total number of entries, average length, maximum length, and number of peptide changes).

Additionally, as we noted a formation of salt bridges between the peptides and LCAT, contact maps between peptide residues 7 and 22, and all LCAT residues were created with a custom script, which simply calculated the number of times the distance between the peptide residue’s backbone bead and LCAT’s backbone beads was under 0.6 nm. The contacts between triplicate simulations were summed, and the maps in the same plots normalized according to the highest number.

### Materials for experiments

ApoA-I mimetic peptides 22A (PVLDLFRELLNELLEALKQKLK), 22A-K22Q (PVLDLFRELLNELLEALKQKLQ) and 22A-R7Q (PVLDLFQELLNELLEALKQKLK) were obtained from Peptide Protein Research Ltd (Fareham, UK). 1,2-dimyristoyl-sn-glycerol-3-phosphocholine (DMPC), 1,2-dipalmitoyl-sn-glycerol-3-phosphocholine (DPPC) and Ergosta-5,7,9(11),22-tetra-3beta-ol (dehydroergosterol, DHE) were purchased from Avanti Polar Lipids Inc (Alabaster, AL). Recombinant human lecithin cholesterol acyltransferase (LCAT) was synthesized by ProSpec-Tany TechnoGene Ltd (East Brunswick, NJ). Cholesterol oxidase (COx) was obtained from Sigma Aldrich (St. Louis, MO).

### Preparation of HDL mimetic nanodiscs for characterization

HDL-mimetic nanodiscs were synthesized as follows. Briefly, phospholipids (DMPC) were dissolved in chloroform (5 mg/mL) in a glass vial, while peptides (22A, 22A-K22Q and 22A-R7Q) were dissolved in 20 mM PBS + 1 mM EDTA (pH 7.4) obtaining a solution 5 mg/mL for each peptide. Phospholipid stock solution was dried with Rotavapor R-200, BUCHI Labortechnik (Flawil, Switzerland) for 1-2 h and then 1 h under nitrogen flow at room temperature to remove the residual organic solvent. The resulting white thin lipid film was rehydrated with 500 μL 20 mM of PBS + 1 mM EDTA (pH 7.4) and vortexed for 5 min. The solution was then homogenized in water bath sonication for 1-2 h at room temperature, obtaining a clear liposome solution. The peptide stock solution was added (2:1 w/w ratio of lipid/peptide). After 2 min of gently mixing, the solution was incubated via heating-cooling cycles (50 °C for 3 min; 4 °C for 3 min) 3 times, obtaining the final HDL-mimetic nanodiscs solution. Hydrodynamic diameters of sHDL were analyzed using dynamic light scattering (DLS) apparatus DLS Zetasizer APS, Malvern Instruments (Westborough, MA). Size distribution profiles are shown as volume intensity average.

### LCAT activity assays

sHDL containing the fluorescent sterol DHE for LCAT activity assay were prepared via Thermal cycling method. In a glass vial, DMPC and DHE in chloroform were combined at a 9:1 molar ratio and were first dried with Rotavapor R-200 at room temperature for 1-2 h and then 1 additional hour under nitrogen flow at room temperature. The thin lipid film was rehydrated with 500 μL of 20 mM PBS + 1 mM EDTA (pH 7.4) and vortex for 5 min. The solution was homogenized in water bath sonication for 1-2h at room temperature. Peptide (22A, 22A-K22A, 22A-R7Q) solution in 20 mM PBS + 1 mM EDTA (pH 7.4) was added obtaining a final lipid to protein w/w ratio of 2:1. After 3 min of a gentle vortex, the solution was incubated via heating-cooling cycles (50 °C for 3 min; 4 °C for 3 min) 3 times. The final concentration of DHE in sHDL-DHE samples was 35 μM. LCAT activity assay was performed in 96-well μL-volume back microplate in triplicate with a final assay volume of 50 μL. LCAT enzyme was dissolved in assay buffer (PBS + 1 mM EDTA + 60 μM albumin; pH 7.4), obtaining a final concentration of 7 μg/mL; sHDL-DHE samples were diluted at different DHE concentrations (0, 10, 20, 35 μM) in assay buffer. 20 μL of sHDL-DHE samples and 20 μL LCAT solution were pre-heated separately at 37 °C for 5 min and then mixed. The plates were incubated at 37 °C in the presence of gentle shaking (250 rpm/h) for 30 min. Reactions were stopped by adding 10 μL of stop solution (PBS + 1 mM EDTA + 5 units/mL cholesterol oxidase COx + 7% Triton X-100). After this, the plates went through a second round of incubation at 37 °C with gentle shaking (250 rpm/h) for 1 h to extinguish the fluorescence of unesterified DHE. After re-equilibrating the plate at room temperature, DHE fluorescence was detected at an excitation wavelength of 325 nm and an emission wavelength of 425 nm using Varioskan LUX, Thermo Scientific. Reactions without LCAT were used for background subtraction, while reactions without LCAT and stop solution lacking COx were used to generate a standard curve for DHE. Reactions were performed in triplicate with two independent experiments. Data were processed via background subtraction (0 μM of DHE) and then divided by the slope of the standard curve, obtaining the amount of DHE-ester resulting in each well. The amount of DHE-ester in each well was divided by the assay time to determine the reaction rate.

### Stability studies

The stability of the samples over time was evaluated by the determination of the hydrodynamic diameter by DLS analysis every 30 days for three months. Samples were stored in glass vials at 4°C protected from the light.

### Electron microscopy imaging

LCAT can hydrolyze lipids even in the absence of cholesterol; for this reason, a short incubation time was used before fixing the solution containing rHDL-LCAT complexes. DPPC was selected as phospholipid because of its lower sensitivity to LCAT hydrolysis with respect to DMPC. sHDL sample (22A+DPPC) for electron microscopy (EM) analysis were prepared by thermal cycling technique, using 20 mM HEPES + 120 mM NaCl + 1 mM EDTA as an aqueous buffer to improve the quality of the images. sHDLs solution was diluted with the same HEPES buffer to obtain a final concentration of 0.10 mg/mL. 10 μg of LCAT were solubilized in 100 μL of 20 mM HEPES + 120 mM NaCl + 1mM EDTA buffer, to obtain a final 0.10 mg/mL enzyme solution. The LCAT– HDL complexes were prepared by pre-heating the LCAT solution (0.10 mg/mL) and rHDLs (0.10 mg/mL) separately at 37 °C for 5 min and then together for 3 min at 37 °C. The sample (3 µl) was allowed to bind on an EM grid (200 mesh Cu + continuous carbon) for 45 s followed by washing three times with ultrapure water and blotting with filter paper before each wash. Staining was performed twice (5 s, then 60 s) with 2 % uranyl acetate, blotting with filter paper before the stain was applied. Finally, the grids were blotted and air-dried. Data were collected using a Hitachi HT7800 electron microscope operated at 120 kV voltage at a magnification of 40,000x, leading to a pixel size of 2.8 Å/pixel. Approximately 1000 micrographs were acquired from sample A, and 164 from sample B with an exposure time of 1 s. Data were processed using RELION (ref) as a part of the Scipion software platform (42) to obtain 2D class averages of the particles.

### Quartz crystal microbalance (QCM)

Before QCM measurements, the LCAT stock solution was prepared by dissolving 10 µg of LCAT in 500 µl of PBS. The solution was centrifuged (14000 g) twice through Amicon Ultra-0.5, 10 kDa cut-off filter (Sigma-Aldrich) for buffer exchange and purification. The retentate was diluted to a total volume of 500 µl (PBS) after each step. Prepared 20 µg/ml (∼430 nM) LCAT solution was stored in −20 °C before further use.

Impedance-based QCM instrument QCM-Z500(KSV Instruments Ltd., Helsinki, Finland) was used for the measurements. The temperature was kept constant at 20 °C. Before measurements, silica-coated sensors (Q-Sense Inc./Biolin Scientific, Västra Frölunda, Sweden) were flushed with 70 % (v/v) ethanol, ultrapure water, dried under nitrogen flow, and oxygen plasma-treated (PDC-002; Harrick Plasma, Ithaca, NY) for 5 min at 29.6 W and 133-173 Pa. Samples were injected into the flow channel using a peristaltic pump system. After the baseline of the signal at different frequency overtones (3, 5, and 7) was stabilized in PBS, a nanodisc surface was formed by flowing nanodisc solution (0.15 mg/ml in PBS, 150 µl/min) on the sensor for 5 min, until the signal was again stabilized. Flow speed was then decreased to 75 µl/min for 5 min flow with PBS to ensure a solid, uniform nanodisc surface. 10 nM (0.47 µg/ml) LCAT was then injected through the flow channel at 75 µl/min for 17 min. Then, the running solution was again changed to PBS to see thefrequency signal stabilize. All measurements were performed three times; in-between measurements, the sensor and the flow channel were cleaned *in situ* by 3 min sequential injections of 20 mM CHAPS, 2 % (v/v) Hellmanex, 70 % (v/v) ethanol, and ultrapure water at 150 µl/min. Frequency overtone signals were normalized, baseline corrected and averaged for each measurement.

## Results and discussion

### The apoA-I mimetic peptides 22A and 22A-K show a different tendency to form antiparallel dimers in the rim of nanodiscs

To investigate the location, conformation, and dynamics of apoA-I mimetic peptides in nanodiscs in silico, we complexed coarse-grained models of DMPC lipids with models of apoA-I mimetic peptides 22A and 22A-K and simulated the systems up to 80 µs, measured in the effective Martini force field time. The proteolytic removal of C-terminal lysine of 22A in serum significantly reduces LCAT activation (6). Thus, we hypothesized that removal of the positively charged amino acid K22 from 22A peptide could affect the average location, orientation, and aggregation tendency of the peptides in nanodiscs, which might give indications behind the decreased LCAT activation potency.

The bulk of the peptides 22A and 22A-K localized in the rim of nanodiscs at a mean radius of 5 nm based on their radial distribution functions (Fig. 1 A), while some individual peptides were observed to transiently travel from the rim to the center of the disc when simulation trajectories were visually inspected (Fig. 1 D). Analysis of the angle distribution between peptides axis and disc normal implied that both peptides are, on average, orientated orthogonally with respect to the nanodisc normal (Fig 1 B). Interestingly, the removal of C-terminal K22 shaped the angle distribution wider compared to 22A indicating that the tendency of the peptides to orient orthogonally was slightly reduced. Then we asked if the amino acid deletion affects the rotational dynamics of peptides. We analyzed rotational autocorrelation functions (ROTACFs) by using the vector formed by N-and C terminal ends of the peptides resulting in faster rotational dynamics for peptide 22A-K when compared to 22A (Fig 1 C). Based on the shapes and half-lives of ROTACFs, we suggest that the peptide dynamics are affected either by the interactions between individual peptides (e.g., formation of rotationally slower dimers in the case of 22A) or peptide-lipid interactions.

**Figure 1.**
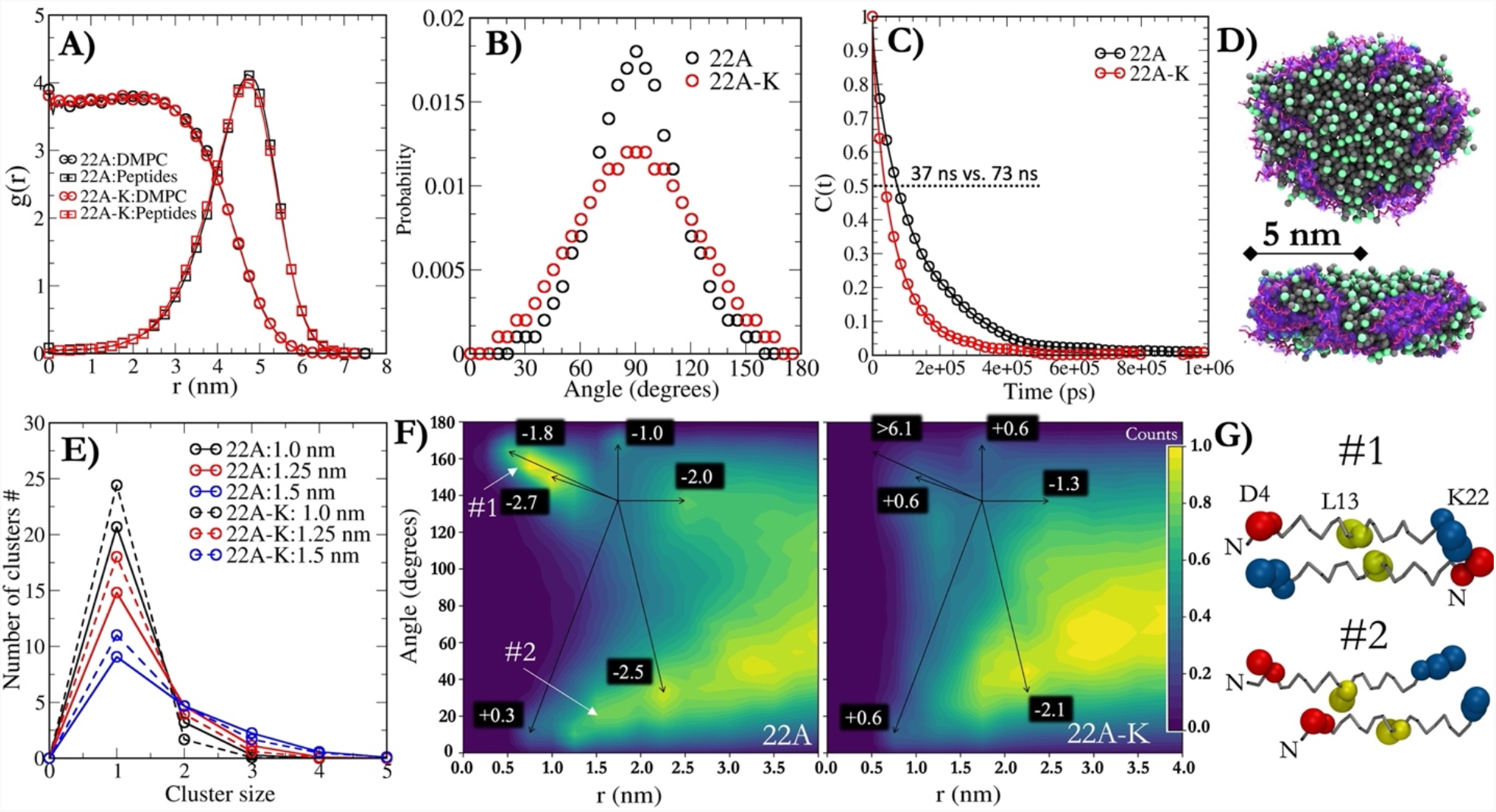
**A)** Radial distribution function profiles for DMPC and peptides in 22A and 22A-K nanodiscs systems. **B)** The angle distribution between axis of the peptides and the normal of nanodisc. **C)** Rotational autocorrelation functions for 22A and 22A-K peptides in the rim nanodiscs. **D)** Simulation snapshots for peptide 22A-DMPC nanodisc showing the molecular organization. DMPC lipids and phosphate group are rendered as grey and green spheres, respectively. Peptides are rendered as purple sticks and transparent surface presentations. Water beads are removed from the figure for clarity. **E)** Number of peptide clusters in 22A and 22A-K systems with different cut-offs. **F)** 2-dimensional contour plots showing the angle distributions between peptide axes as a function of distance in 22A and 22A-K systems. The white numbers on the black squares show the free-energy differences (kJ/mol) to the reference position of 1.75 nm and 135° **G)** Representative snapshots for 22A peptide conformations at #1 and #2. Backbone beads of peptides are rendered as grey sticks. Blue, red, and yellow beads represent K22, D4 and L13 amino acids.

Next, our target was to investigate the dimerization and oligomerization tendencies of 22A and 22A-K. For this purpose, a cluster analysis was carried out to find the number of peptide clusters and their relative probabilities during the simulations (Fig 1 E). The 22A and 22A-K peptides preferred to be mainly in a monomer form when a cut-off of either 1.0 nm or 1.25 nm was used. Also oligomerized peptides were present with either cut-off used, but the aggregation tendency was less pronounced in the case of 22A-K (lower number of monomers). Specifically, depending on the cut-off used, 25-68% and 14-61% of the total number of 22A peptides 22A and 22A-K were present in oligomers, respectively. We sought to explain this different preference for oligomer formation by exploring the orientational behavior of neighboring peptides as a function of residue L13 distance (Fig1 F). Intriguingly, the analysis revealed that 22A peptides had a strikingly different angle-distance behavior when compared to 22A-K since 22A peptides adopted relatively restricted antiparallel orientation with respect to each other when the distance of the neighboring peptides is 0.5-1.5 nm. The preferred angle between two 22A peptides was ∼150-160 degrees when the distance was ∼0.5-1.0 nm (antiparallel conformation #1, Fig 1 G). 22A peptides were also able to form parallel dimers, but the distance separation in this conformation was slightly increased since the peptides are imperfectly aligned when the peptide sequences are considered (parallel orientation #2, Fig 1 G). It is clear from the 22A-K profile that the antiparallel orientation does not take place, although there are very weak probability densities present near 0.5-1.0 nm. In addition, the angle probabilities at the distance range of ∼0.7-1.5 nm are the same across the different angle values. Moreover, the overall shapes of angle-distance profiles were drastically different between peptides 22A and 22A-K. We also determined the free-energy changes with respect to the free-energy maximum seen in the angle-distance profile of 22A (1.75 nm and 135 degrees) to estimate the thermodynamic stability of dimers. The antiparallel orientation was favored by 3.3-7.9 kJ/mol or more in the case of 22A when compared to 22A-K. However, the antiparallel dimers in the case of 22A were thermally relatively short-lived as the free-energy barrier associated with their monomerization is only −2.7 kJ/mol, which was also visible in the simulation trajectories.

To conclude, removing the C-terminal lysine of 22A strikingly alter the dynamics and dimerization properties of apoA-I mimetic peptides studied here. These features might play an essential role in the activation of LCAT since the correct registry of two antiparallel apoA-I proteins is crucial for efficient LCAT activity (33). The proper registry of apoA-I proteins, and the antiparallel dimers of 22A peptides shown here, might generate an active conformational subunit for efficient LCAT binding and activation. Moreover, it has been suggested that the specific LCAT-interacting motif generated by two apoA-I monomers might facilitate the entry of free CHOL to the active site of LCAT through an amphipathic “presentation” tunnel(43).

### Lecithin-cholesterol acyltransferase localizes into the rim of nanodiscs and adopts spatially restricted location and conformation when the lid is in the open state

To investigate how LCAT is localized and oriented in the nanodiscs and if these features are influenced by removing the end lysine from 22A, we placed LCAT into the peptide-DMPC nanodiscs and carried out CG-simulations for systems 22A and 22A-K. In both cases, inactive (the lid closed) and active (the lid open) forms were investigated (Fig 2A). We anticipated that LCAT might end up in semi-stable states as it travels on the disc and interacts with peptides. Therefore, triplicate simulations (3 × 40 µs in effective Martini time) were run for each configuration to eliminate this possibility.

**Figure 2.**
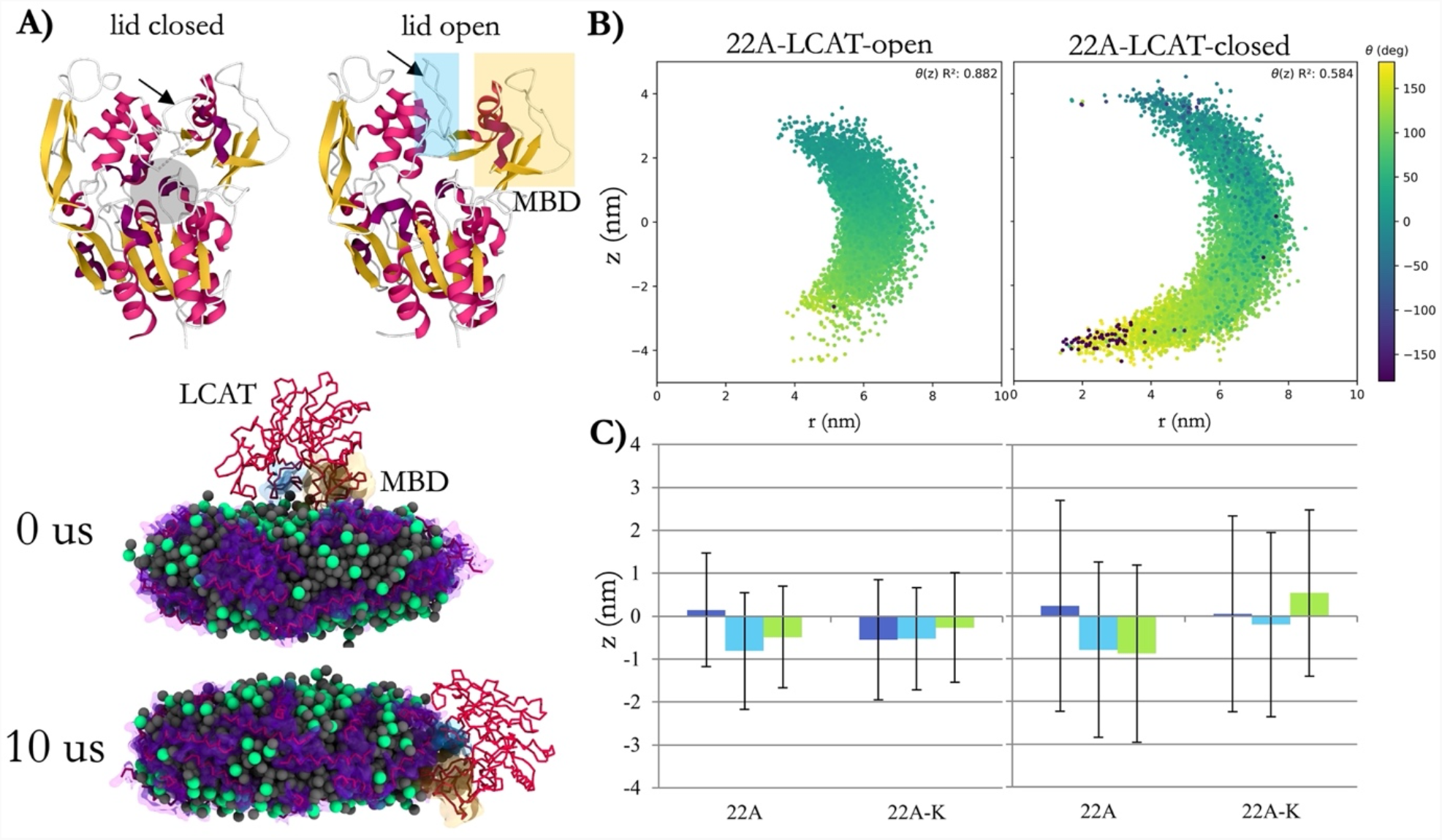
**A) TOP:** X-ray structures of open (6MVD) and closed (4XWG) LCAT showing the locations of membrane binding domain (yellow box) and lid (blue box). The grey and transparent sphere marks the location of the active site. **BOTTOM:** Snapshots of open LCAT-22A simulation system from the start (0 us) and end (10 us) of simulation showing the relocation of LCAT to the edge of the nanodisc. **B)** The z-axis location and orientation of LCAT with respect to disc normal in the rim of nanodisc (angle distribution analysis with respect to nanodisc plane or normal) as function of distance from the geometrical center of the disc. The correlations between z-axis location and angle are shown in top right corner of the profiles. **C)** The distance averages of LCAT with respect to z-axis (parallel to disc normal) for each triplicate simulation of 22A and 22A-K systems with open (left) or closed (right) LCAT. The errors bars are standard deviations.

First, we examined the propensity of LCAT to stay attached to the nanodiscs by measuring its contacts with the peptide beads. In Martini force field time, one simulation with 22A-LCAT-open and two with 22A-LCAT-closed took 8 µs to find equilibrium, whereas the rest of the 22A and 22A-K systems took 4 µs. In all systems, LCAT positioned itself on the disc perimeter with its active site cleft pointing to the lipid interface (Fig 2A). No detachments of LCAT from the surface of nanodiscs occurred. Once the systems reached spatial equilibrium, the spatial position and orientation of LCAT respective to the disc were characterized by measuring its distance along with the disc from the lipid center of mass (r), its position along the disc normal (z), and its pitch angle, θ, to the disc normal (Fig 2B). Interestingly, the z position of LCAT is highly correlated to its angle, meaning that as LCAT moves above and below the disc at the perimeter, its active site remains covered by pointing to the lipid interface, even when the lid is closed. Conversely, LCAT in its active form with the lid open is conformationally and dynamically more restricted at the perimeter of the disc, as can be seen in the standard deviations which are doubled with LCAT closed form. Similarly, in the open form, LCAT generally stays slightly below the disc, and due to the high correlation, with a well-defined orientation. The location and orientation of LCAT in the rim of nanodisc were unaffected by the activity reducing peptide mutation in 22A-K (Figure S1 and Table S1).

Next, we analyzed the solvent-accessible surface areas (SASAs) of all LCAT residues in the water phase and when bound to nanodiscs to scrutinize our simulation LCAT model against previously published experimental hydrogen-deuterium exchange (HDX) data (34). Both open and closed forms of LCAT were analyzed. The results acquired in the water surrounding were subtracted by the values obtained when LCAT was bound to nanodisc to produce data highlighting the LCAT regions interacting mainly with peptides or lipids in the nanodisc (Fig 3 A). In this analysis, LCAT chiefly utilized the amino acids in the MBD (40-70), αA-αA’ loop (110-130), and lid loop regions when attached to the nanodisc. However, weak interaction sites were also registered in two areas spanning amino acids of 330-340 and 370-390. These results are in line with the HDX data of Manthei et al. when LCAT was bound to HDL particles comprised of two apoA-Is (34). That is, the MBD, αA-αA’ loop, and the lid regions were the main sites that were protected from HDX when LCAT was attached to HDL particles (the experimentally determined shielded areas are shown with blue boxes in Fig 3B). It is worth noting that our simulation system is based on short and dynamic apoA-I mimetic peptides in contrast to the more spatially constrained full-length apoA-Is, a fact that could explain the differences seen in the profile. In addition, in our previous work, we revealed utilizing atomistic MD simulations that the hydrophobic amino acids present in these regions were considerably less accessible to water when the closed-form of LCAT in water was compared to the open-form of LCAT that was bound to a lipid bilayer (44).

**Figure 3.**
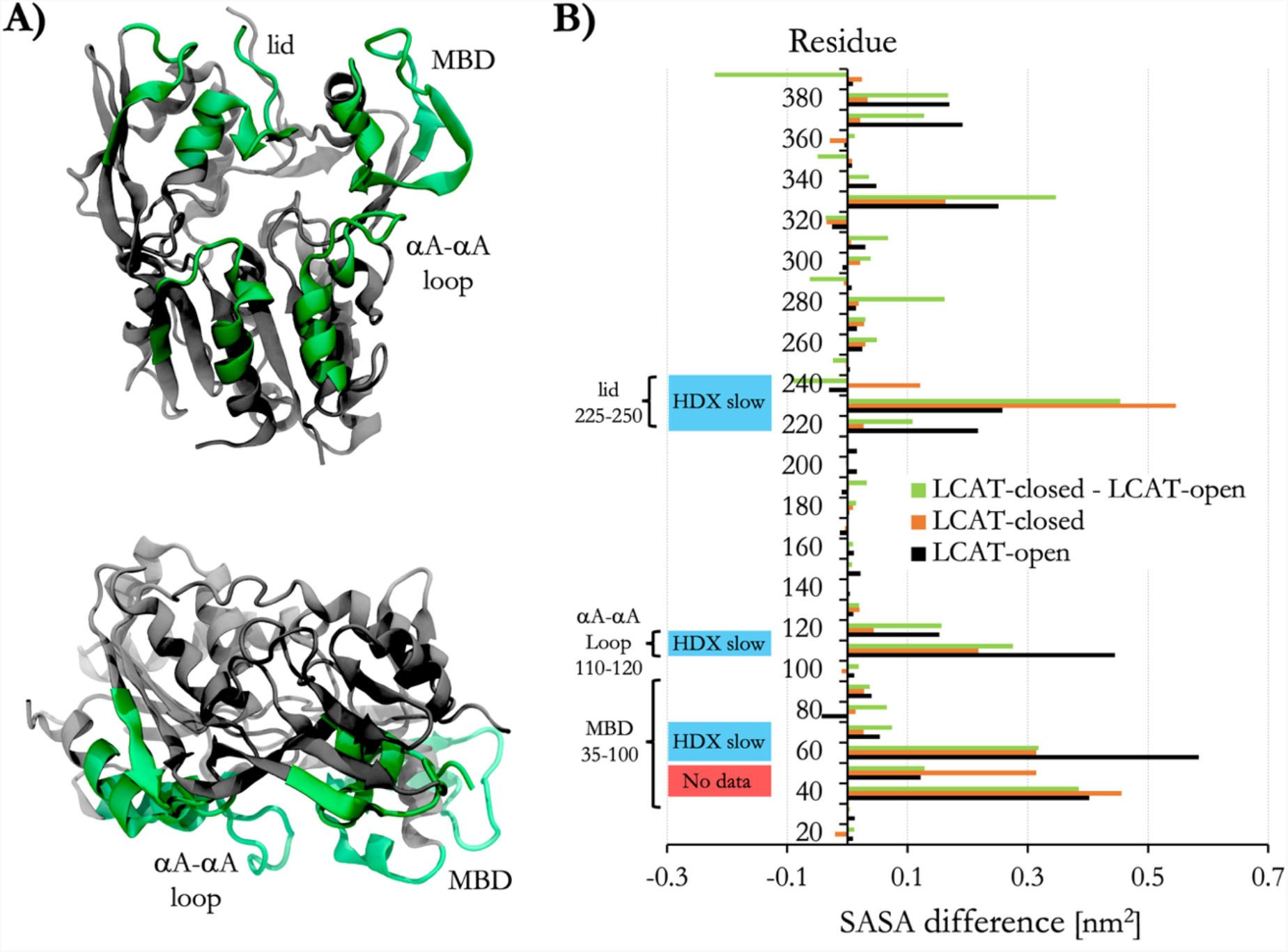
**A)** Color-mapped X-ray structures of open LCAT (6MVD) showing the structural regions of LCAT that are less accessible by water (green) when LCAT is attached to nanodiscs in simulations. **B)** Solvent accessible surface area (SASA) analysis showing the differences in water exposures as a function of amino acid sequence of LCAT. Blue boxes show the protected regions based on previously determined hydrogen-deuterium exchange (HDX) data when LCAT attaches to HDL particles (34). Red box indicates the region for which no HDX data was available.

Since the localization and orientation of LCAT were spatially restricted, especially in the open form, in our simulations, we sought to validate this experimentally. We used negative staining EM to verify our results regarding the location and orientation of LCAT in the rim of peptide nanodiscs. Fig 4A shows representative negative staining EM images from 22A nanodiscs with and without LCAT. The average size of 22A nanodiscs in the absence of LCAT was ∼10 nm with a peptide-lipid ratio of 1:7 (Fig 4A, inset). Image classification and 2D averaging revealed populations of nanodiscs where LCAT was either unbound, or where one copy of LCAT located in the rim of nanodiscs (Fig 4 B and C). Surprisingly, we also found a minor subpopulation where two putative LCAT enzymes were bound to nanodiscs with a well-defined distance between the two enzymes (Fig 4 C). The EM images indicate that LCAT prefers to locate on the rim of nanodiscs, and it is conformationally restricted enough to produce image subpopulations showing a seemingly similar orientation of LCAT with respect to the edge of nanodiscs (Fig 4C). As the dimensions of LCAT in the rim of nanodiscs were similar between image subpopulations, we extracted the average orientational coordinates of LCAT in its open form from our CG simulations based on the distance and angle calculations shown in Fig 2B and C. Subsequently, the average representative structure of LCAT bound to 22A nanodisc was 2D-fitted with EM derived images so that the normal of nanodiscs were pointing towards the same direction in MD and EM derived images. As shown in Fig 4D, the location and dimensions of the simulation-derived average LCAT structure closely matched the dimensions and shape of LCAT in a 2D class average from EM, adding support to our simulation results concerning the location and orientation of LCAT.

**Figure 4.**
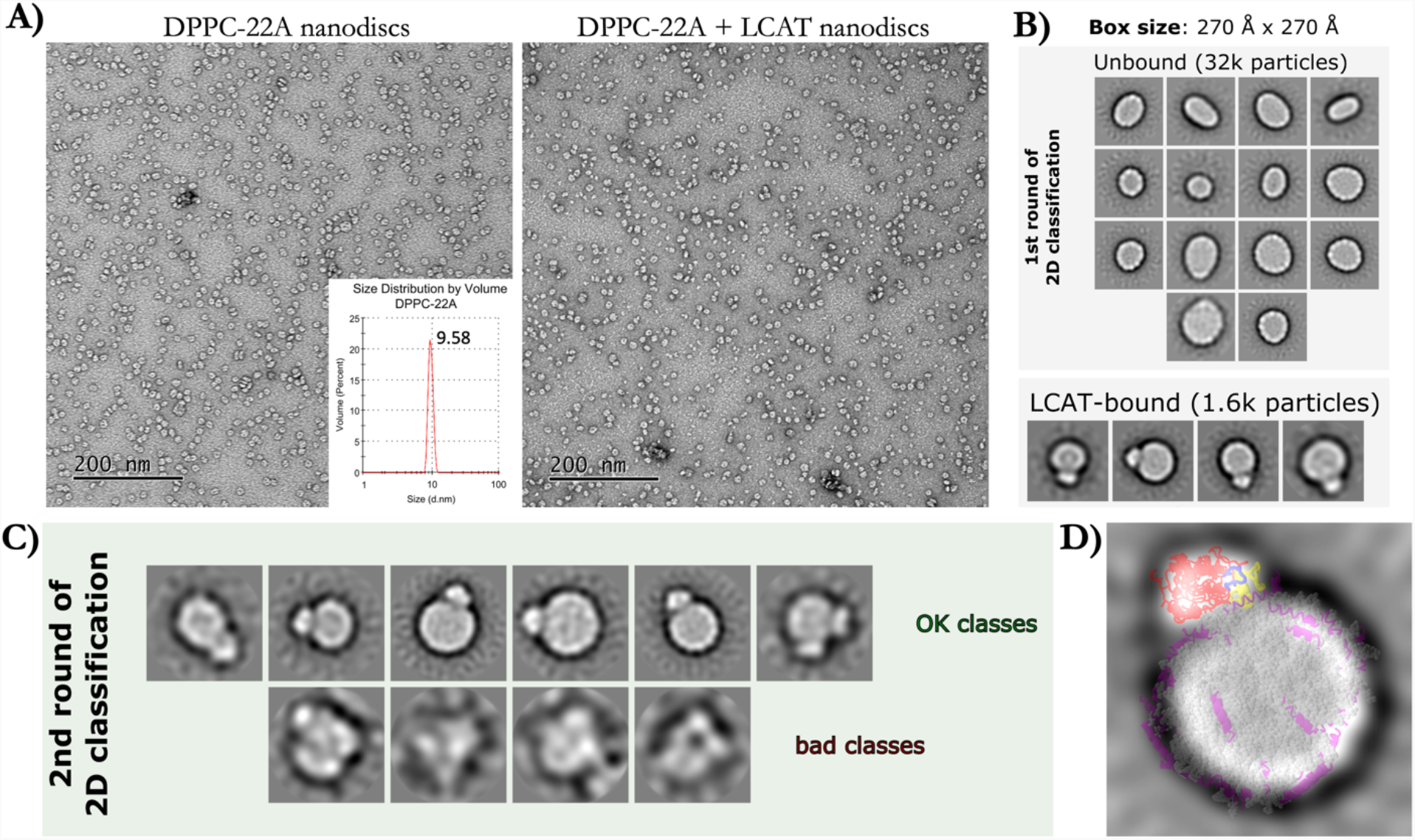
**A)** The negative staining electron microscopy images of DPPC-22A nanodiscs without (left) and with (right) LCAT. The inset in the image without LCAT shows the particle sizes (22A-DPPC) determined by dynamic light scattering before introducing LCAT. **B)** Two-dimensional class averages of nanodiscs with LCAT and DPPC-22A. The top image shows class averages of discs without LCAT and the bottom one shows class averages for discs with LCAT bound to the edge of nanodiscs. **C)** Class averages after further classifying the LCAT-bound nanodiscs, showing good and bad classes. **D)** An overlay of EM LCAT-disc image and the MD simulation-derived coordinates resembling the appearance of LCAT in the rim of nanodisc based on the location and conformational analysis shown in Fig 2.

Taken together, our data indicate that LCAT prefers to be localized in the rim of nanodiscs, similar to the case of HDL particles with apoA-I (34). The regions that LCAT utilizes in the attachment to the nanodisc’s surface agree with the previous experiments and MD simulations (44–47). Surprisingly, we observed that open LCAT adopts a more restricted orientation spatially with respect to nanodisc normal and surface when compared to the closed form. This orientational correlation matched well with the negative staining images. The reason for the more restricted orientation of LCAT in the open state might be a more specific and stronger interaction with the apoA-I mimetic peptides.

### ApoA-I mimetic peptides bind specifically to the lid-MBD groove of open LCAT and subsequently form transient antiparallel dimers

As the apoA-I mimetics act as cofactors in LCAT-mediated CHOL esterification, we sought to characterize the main binding and interaction site for 22A peptides in the structure of LCAT and how this interaction is modulated by the removal of the C-terminal lysine of 22A. To achieve this, we calculated the spatial densities of peptides 22A and 22A-K around the open and closed forms of LCAT. The analysis indicated that 22A peptides concentrated next to the lid and MBD regions of LCAT when LCAT was in the closed state (Fig 5A left), whereas in the case of open LCAT, peptides showed a strong preference to bind and interact with the groove formed by the lid and MBD of LCAT (Fig 5A right, site A). Another, albeit slightly less prominent, spatial density hot spot was registered in the vicinity of the active site tunnel opening and the αA-αA’ loop. However, this hotspot was less consistent across the different simulations when compared to site A. The spatial density analysis showed similar results for 22A-K. Next, we calculated the temporal occupancies of peptides 22A and 22A-K in site A to estimate their relative binding strengths that may explain their different potencies to activate LCAT. As shown in Figure 5B, the dwelling time of 22A-K in site A was markedly lower when compared to 22A, and the behavior was the same across all replicate simulations carried out. This difference seems to be caused by 22A both having a higher tendency to enter the site and a longer residence time there (Table S2).

**Figure 5.**
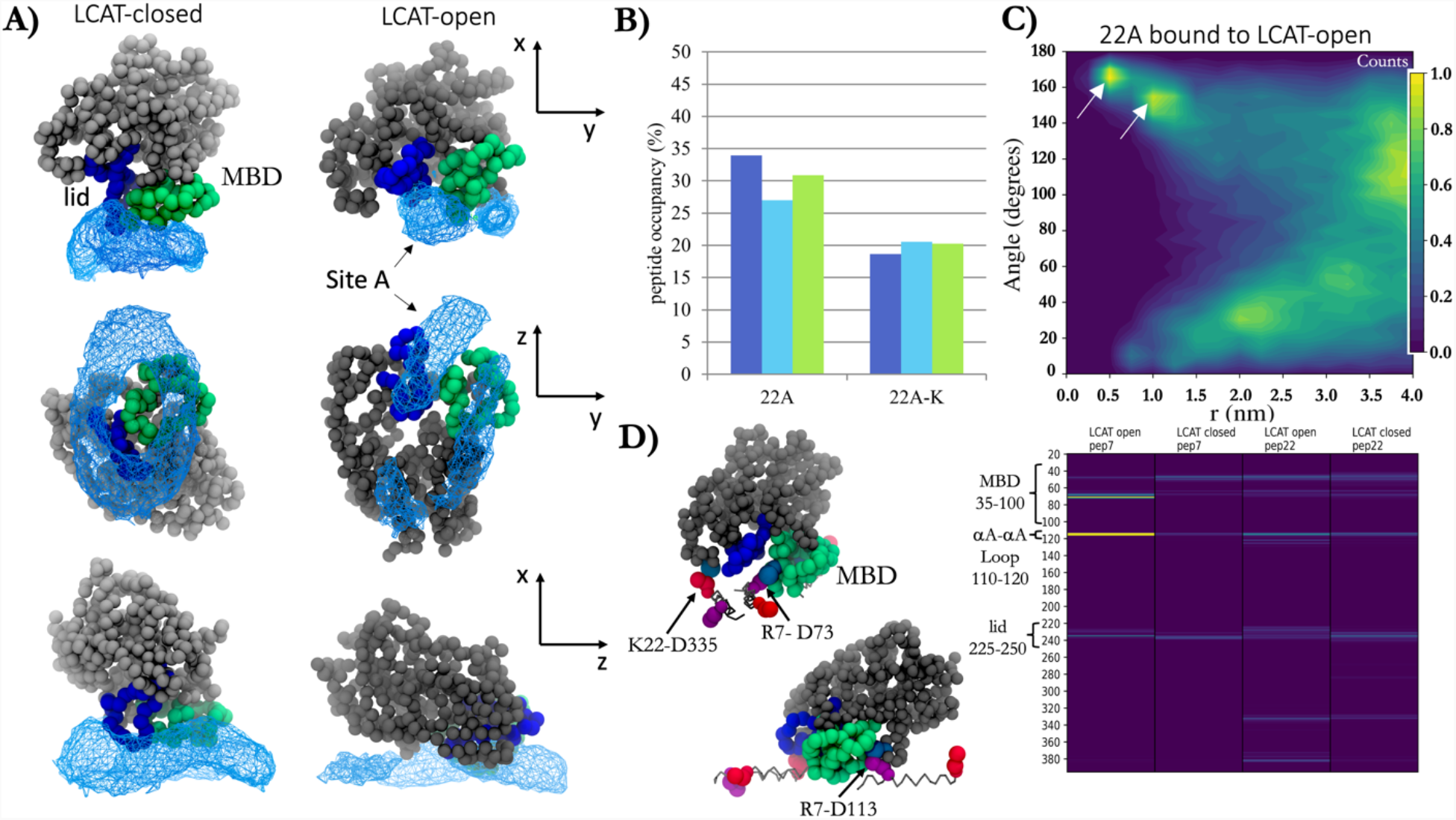
**A)** Spatial probability densities for 22A peptides next to the closed and open LCAT. LCAT is rendered using grey, green (MBD), and blue (lid) spheres. The spatial density of 22A around LCAT is shown as a light blue wireframe presentation. **B)** The temporal occupancy percentages of peptides 22A and 22A-K in the lid cavity. 22A **C)** The angle between peptides 22A and 22A-K as a function of distance when one of the peptides is bound to the lid-MBD groove. White arrows point the antiparallel dimer population maxima. **D) LEFT:** Visualization of major salt-bridges formed between 22A peptides and LCAT. Grey sticks are drawn between backbone beads of peptide 22A. R7 and K22 residues are shown as purple and red spheres, respectively, whereas ASP residues of LCAT are light blue. **RIGHT:** The salt-bridge contact heat maps between LCAT and residues R7 and K22 of peptide 22A.

We further explored how the binding of peptides to the interaction site affects their ability to form the antiparallel dimers since this might be an essential prerequisite for the apoA-I mimetic peptides when the activation of LCAT is considered. Strikingly, 22A peptides were still able to form antiparallel peptide dimers when one of the peptides was bound to LCAT simultaneously (Fig 5C). Namely, the neighboring amphiphilic helixes could adopt an antiparallel conformation at the distance of 0.5-1.0 nm and angle varying approximately between 160-170°. Consistently, the antiparallel dimerization was almost completely absent in the case of 22A-K (Figure S2).

To characterize the interactions between 22A and LCAT, we calculated the number of contacts between different amino acids of 22A and LCAT. We found that R7 and K22 formed salt bridges with the MBD, αA-αA’ loop, and the lid regions of LCAT (Fig 5D, right). Through visual inspection of the simulation trajectories, we observed that R7Q of 22A interacted prominently with the negatively charged amino acid in the αA-αA’ loop (D113) when the lid was in the open state (Fig 5D, left). Additionally, both amino acids formed salt bridges with a specific negatively charged amino acid in the lid-MBD cavity groove (D73) in the case of both open and closed LCAT. In addition, in the case of 22A, the antiparallelly dimerized peptide was able to form specific salt bridges with residue D335 of LCAT.

Our results highlight that there is a specific binding site for apoA-I mimetic peptides in the open LCAT structure which may be a prerequisite for CHOL esterification. Peptide 22A was able to form antiparallel dimers while in site A, suggesting that this would also be the primary binding site for apoA-I. When comparing the interactions between LCAT and peptide 22A dimer to the 3D reconstitution of LCAT with respect to two apoA-Is in native HDL particles, it is clear that the interaction modes are similar (34). Intriguingly, this indicates that even short apoA-I mimetic peptides can arrange themselves next to LCAT in a manner that resembles the configuration of two apoA-Is. This kind of orientation may mitigate the entry of CHOL into the active site between amphiphilic α-helixes. However, since the residence time of peptides 22A and 22A-K in site A was ∼30 % and ∼20% of the total simulation times, the free-energy of binding of both peptides to site A is slightly positive. Therefore, it might be interesting to experiment if an apoA-I mimetic peptide variant with a negative free-energy of binding can considerably increase LCAT activity or not.

### Apolipoprotein A-I mimetic peptide occupancies in the lid groove of LCAT correlate with their activation potencies

Based on the simulation results above, we designed two 22A variants (22A-R7Q and 22A-K22Q) to study the effect of charge neutralization of residues R7 and K22 on the activation of LCAT. For this purpose, we reconstituted peptide-DMPC nanodiscs with the fluorescent sterol DHE to measure their LCAT activation potencies. Both peptide variants 22A-R7Q and 22A-K22Q were able to form peptide-DMPC nanodiscs with and without DHE (Fig 6A). The sizes of nanodiscs (9.1-9.9 nm vs. 9.6-10.3 nm) did not change upon incorporating DHEs into the nanodiscs, as shown in Fig 6A. As the different lipid-protein ratios and the total charge of the peptides might affect nanodisc stability, e.g., through the change of interfacial tensions, over time, we also monitored the stability of nanodiscs up to 90 days (Fig 6 B). We did not observe any differences in stabilities between the two nanodisc types, which indicates similar biophysical properties of the nanodisc surfaces when the stability is concerned.

**Figure 6.**
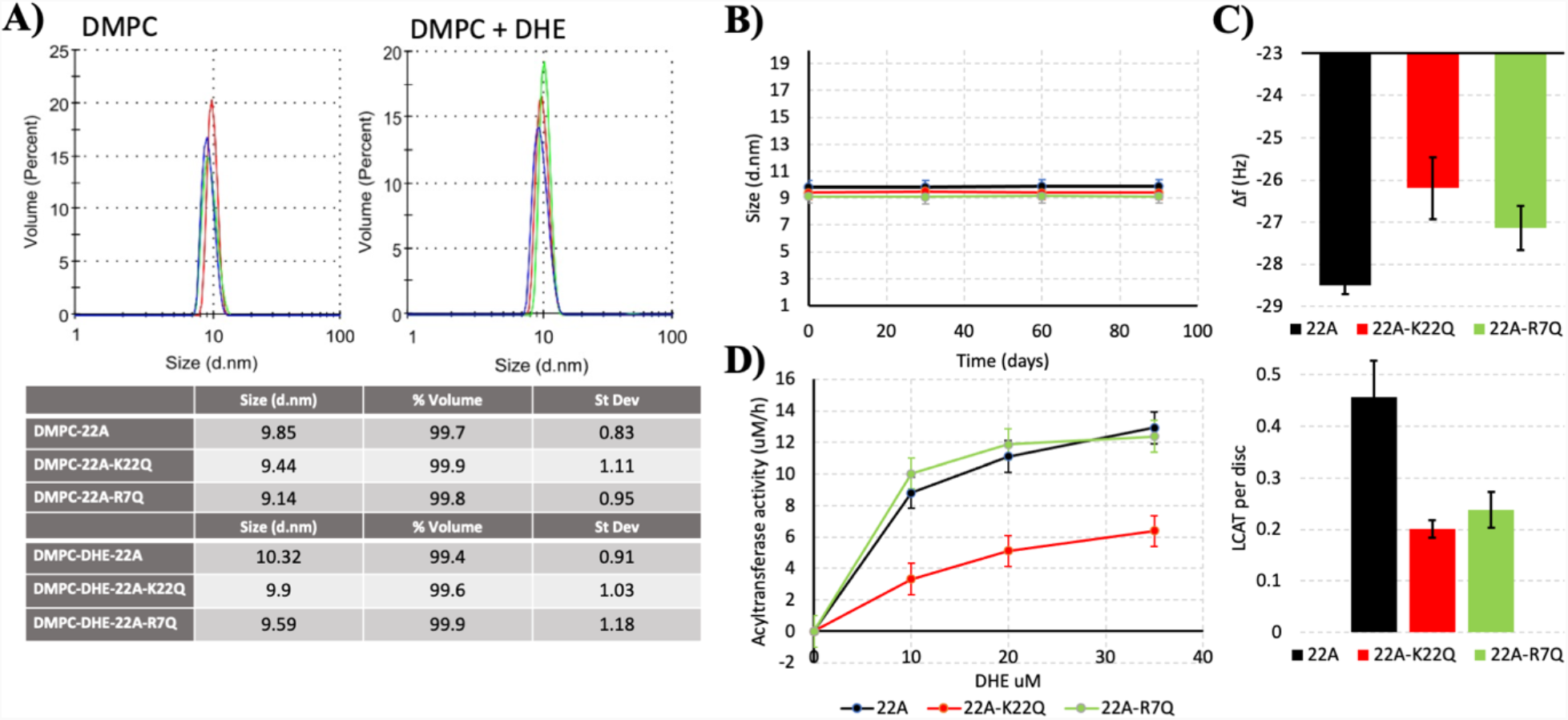
**A)** The effect of peptide sequence on the average size of nanodiscs with and without dehydroergosterol. **B)** Stability of nanodiscs up to 90 days. **C) Top:** Quartz-crystal microbalance results for the binding of 22A, 22A-K22Q, and 22A-R7Q nanodiscs to the sensor surface. **Bottom:** Estimated number of LCAT enzymes bound per disc based on adsorbed mass on the sensor after LCAT injection. **D)** Acyltransferase activity of LCAT with different DMPC/DHE/peptide nanodiscs

Next, to assess the effect of the total negative charge of peptides (22A-R7Q and 22A-K22Q vs. 22A) on LCAT binding to nanodiscs, we utilized the QCM technique to monitor the mass of LCAT adsorbed on peptide-lipid nanodiscs that were spread on the QCM sensor. The results in Fig 6C indicate that the amount of nanodiscs on the sensor after spreading depends on the peptide variant complexed with lipids (Fig 6C, top). That is, negatively charged peptide variants resulted in decreased adsorption of nanodiscs on the sensor. The negatively charged peptides also reduced the adsorption of LCAT on the discs (Fig 6 C bottom), and our calculations suggest that approximately one LCAT was bound to every two nanodiscs in the case of 22A, and one LCAT was attached to every five nanodiscs in the case of 22A-K22Q and 22A-R7Q. LCAT activity assays indicate that variant 22A-R7Q showed nearly identical activity with 22A, but variant 22A-K22Q decreased the LCAT activity by 50%, which agrees well with previous measurements with the 22A-K variant (21A) (6).

As 22A-K22Q decreased the LCAT activation potency, whereas 22A-R7Q did not, we simulated both variants to inspect if their organization and interaction with LCAT differ similarly as in the case of 22A and 22A-K. Although not statistically relevant, our QCM results mirrored the closed LCAT simulations, where LCAT dissociated at 32 µs un two of the three K22Q simulations and at 24 µs in one of the R7Q simulations in effective Martini time. With open LCAT both peptide variants were able to bind to site A and form antiparallel dimers equally well compared to 22A (Fig 7A, Fig 5C and Fig S3). The location and orientation of LCAT were also similar as in the case of 22A and 22A-K (Figure S4 and Table S1). When the occupancies of peptide variants in the site were calculated, it was found that 22A-K22Q had a reduced preference to occupy site A when compared to 22A-R7Q and 22A (Fig 7B and Table S2). The contact heat map analysis of 22A-R7Q show that the neutralizing R7 considerably reduces its interaction with αA-αA’ loop amino acid D113 (Figure S5). However, this did not have an effect to the activity of LCAT indicating that it is not a crucial interaction site regarding the esterification of CHOL molecules.

**Figure 7.**
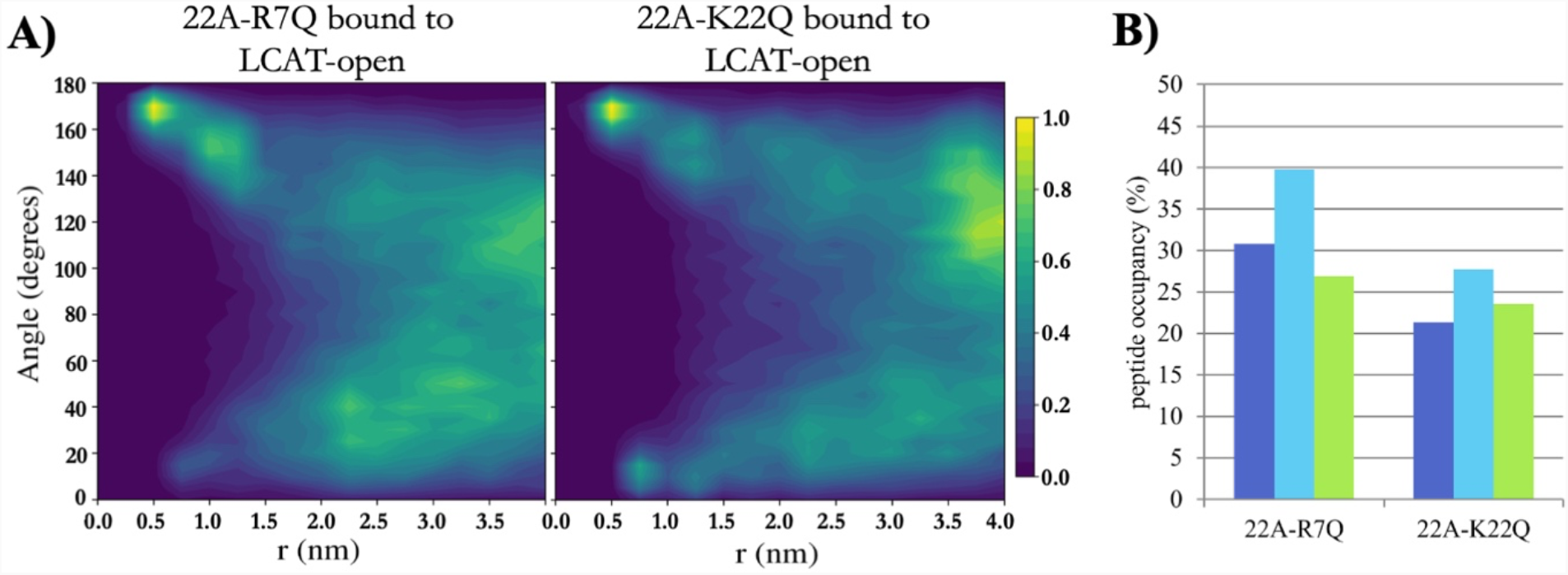
**A)** 2-dimensional contour plot showing the angle between peptides 22A-R7Q and 22A-K22Q as a function of distance when one of the peptides is bound to the lid-MBD groove. **B)** The temporal occupancy percentages of peptides 22A-R7Q and 22A-K22Q in the lid-MBD groove.

Our results suggest that the occupancy of apoA-I mimetic peptides in the lid groove correlates with their ability to promote CHOL esterification. In addition to this, the ability of peptides to form dimers when bound to the interaction site might play an important role in the activation. The binding of peptides into the binding site and the subsequent formation of antiparallel dimers might generate a thermodynamically or kinetically favored pathway for the CHOL into the active site after lipolysis of PLs. This hypothesis further agrees with the research carried out and shows that a correct apoA-I registry is a requirement for increased LCAT activity (33). The importance of positively charged amino acid R127 was also shown to be necessary for LCAT esterification action, and we argue that this might be because of the lower occupancy of correct apoA-I helixes in the lid-MBD groove of LCAT (32). Previously, we have shown that the entry of free CHOL into the active site of LCAT was rendered highly favorable after the acyl intermediate of LCAT is formed, and no involvement of apoA-I or apoA-I peptides is needed (44). However, we used planar lipid bilayers to show this behavior, and it might well be that more assistance is needed for the entry of CHOL in the LCAT. This is because the LCAT action presumably takes place in the rim of the nanodisc, which is highly positively curved (Fig 2A), which might reduce the localization of free CHOL to the edge of the discs. Interestingly, we have previously shown that free CHOL molecules concentrate next to apoA-I monomers in spherical HDL-like lipid droplets (48–50). Further, previous atomistic and coarse-grained MD simulations have suggested that the helix X can facilitate the entry CEs into the hydrophobic tunnel of cholesterol ester transfer protein (51). A similar kind of guiding mechanism might also play a role in the case of apoA-I mimetic peptides when they are bound to the lid residing groove. Namely, apoA-I mimetics bind and orient themselves in the lid-MBD groove of LCAT to enable peptide-harbored CHOL to enter the active site of LCAT in a highly positively curved PL surface. Interestingly, the experimental evidence points out that a set of positive allosteric modulators can facilitate LCAT activity when 22A-based nanodiscs are used in the enzymatic assays (52). How the allosteric modulators boost the activity with 22A peptides remains unknown. Our previous modeling evidence suggested that the positive allosteric modulators change the orientation and spatial free-energy profile of MBD, which might facilitate the opening of the lid, generating a higher population of LCAT enzymes in the open state (53). However, in the light of this study, the orientational shift of MBD induced by the drugs might also facilitate the binding of apoA-I peptides to the lid groove or render the active site tunnel opening more accessible for CHOL molecules.

## Conclusions

Here, we sought to learn how apoA-I mimetic peptides interact with LCAT and how the interaction features correlate with LCAT activities in nanodiscs. Our simulation results agreed well with existing experimental evidence regarding what enzyme regions LCAT utilizes in the attachment to nanodiscs. In addition, the simulation-derived average location and orientation of LCAT in the rim of nanodisc matched the negative staining EM images produced here, further validating the simulation models employed in the current study. Interestingly, our simulation results highlight the different tendencies of 22A peptides and their variants to form antiparallel dimers when bound to site A in LCAT. This behavior, however, does not entirely explain the different activities as variant 22A-K22Q also formed antiparallel dimers equally well compared to 22A but lowered the activation of LCAT by 50%. Another factor that could play a part in the LCAT activation is the binding strength of apoA-I mimetics into the lid-MBD groove. Namely, our peptide occupancy results derived from simulations strongly correlated with the LCAT activities and, thus, we propose that the interaction of apoA-I mimetics with this site is crucial for the LCAT activation, and the binding strength together with the antiparallel dimerization tendencies of the mimetics can modulate the activity of LCAT. Further investigations are needed, for example, to elucidate how apoA-I mimetic peptides while residing in the lid-MBD cavity promote CHOL esterification and how important role the dimerization plays in the end. In toto, the results, mechanistic insights, and methodology reported here will provide a blueprint that can be utilized to design novel drug formulations against LCAT deficiencies and cardiovascular diseases in the future.

## Acknowledgments

The authors wish to acknowledge CSC – IT Center for Science, Finland, for computational resources and the Electron Microscopy Unit of the Institute of Biotechnology, University of Helsinki, for access to facilities. This research received funding from the Academy of Finland (298863 to A.K.)

## Notes

### Competing Interest Statement

The authors have declared no competing interest.

