## Supplementary Information for "Mechanistic insights into the activation of lecithin-cholesterol acyltransferase in therapeutic nanodiscs composed of apolipoprotein A-I mimetic peptides and phospholipids"

**Figure S1.** Position and orientation of LCAT relative to 22A or 22A-K nanodisc normal based on triplicate 10 µs simulations with LCAT.

**Figure S2.** Bound 22A-K peptide's angle as a function of distance to other peptides based on extended triplicate 20 µs LCAT open simulations.

**Figure S3.** Angle as a function of distance between all peptide pairs based on 20 µs simulations without LCAT. Free energy differences to reference position of 1.75 nm and 135° are shown.

**Figure S4.** Position and orientation of LCAT relative to 22A-R7Q or 22A-K22Q nanodisc normal based on triplicate 10 µs simulations with LCAT.

**Figure S5.** Contact heat map between peptide residue and all LCAT residues.

**Table S1.** Peptide orientation when bound based on triplicate 10 µs LCAT open simulations. Data as mean (SD of means).

**Table S2.** Supplementary data on peptide occupancy based on extended triplicate 20 µs LCAT open simulations. Data as mean (SD of means).

#### Figure S1

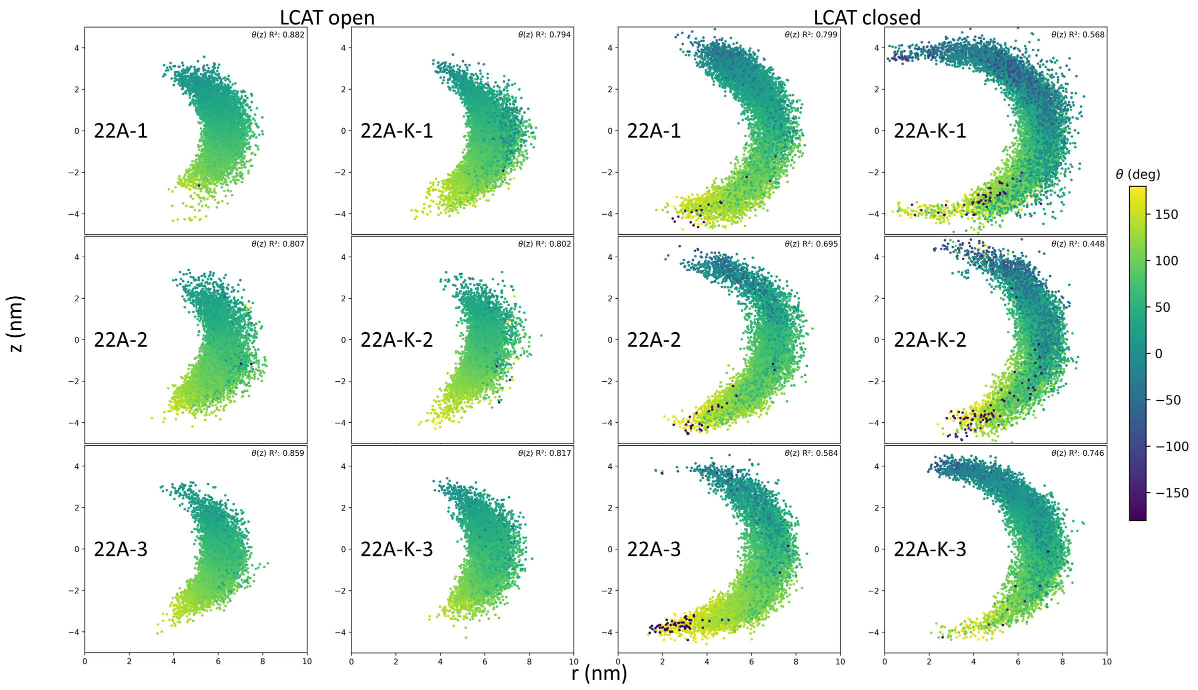


#### Figure S2

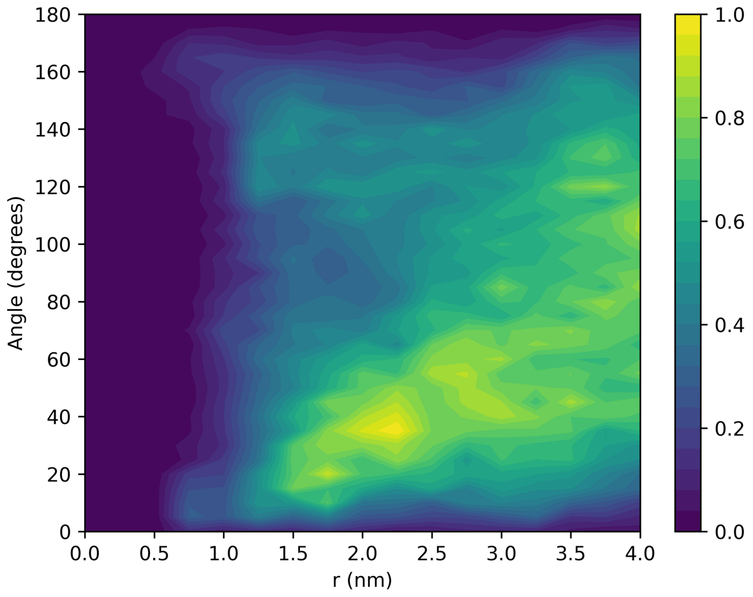

### Figure S3

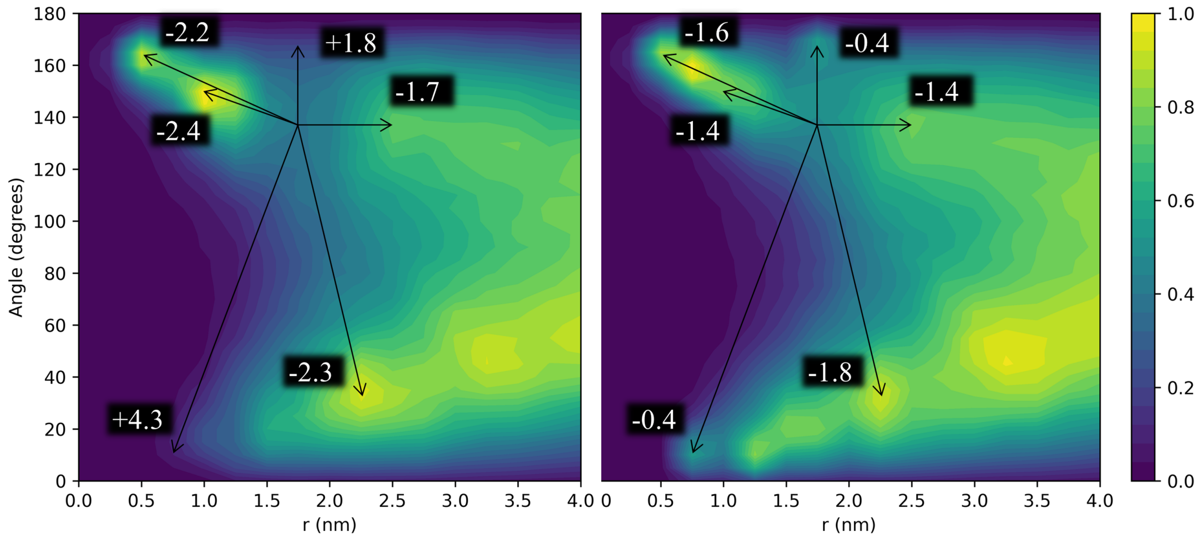


### Figure S4

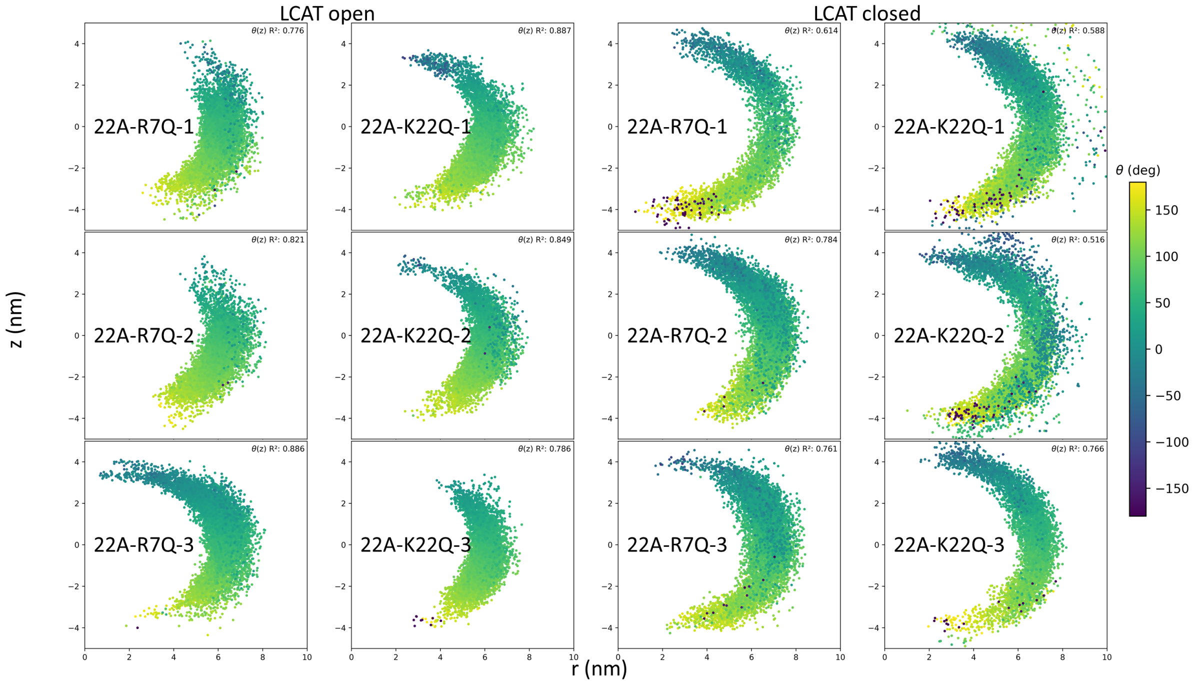

#### Figure S5

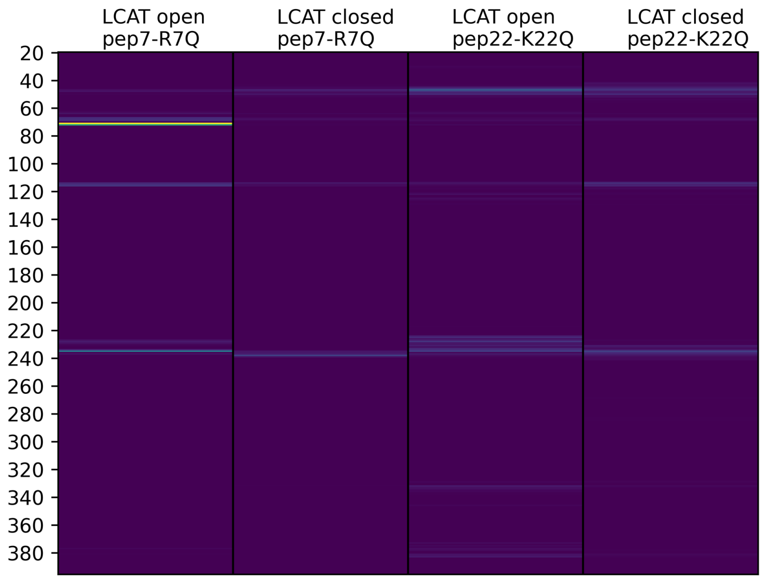


#### Table S1

| Peptide | Peptide vs. CYS31-GLY308 angle (degrees) | Peptide spin (degrees) |
|---|---|---|
| 22A | 45.2(0.8) | 2.8(2.8) |
| 22A-K | 47.5(0.9) | 8.8(1.7) |
| 22A-R7Q | 48.7(2.5) | 11.4(3.4) |
| 22A-K22Q | 43.9(1.1) | 4.2(1.0) |

#### Table S2

| Peptide | Occupancy (%) | Mean occupancy length (ns) | Max occupancy length (ns) | Num of entry events (n) | Num of peptide changes (n) |
|---|---|---|---|---|---|
| 22A | 31.5(1.8) | 4.0(0.1) | 83.7(11.4) | 1463(118) | 6(2.2) |
| 22A-K | 17.9(3.2) | 3.0(0.1) | 42.0(5.9) | 1144(165) | 10(0.0) |
| 22A-R7Q | 32.5(5.4) | 4.1(0.5) | 71.0(19.4) | 1485(85) | 5(0.8) |
| 22A-K22Q | 24.2(2.6) | 3.8(0.3) | 62.3(13.6) | 1206(188) | 5(0.8) |
